# Experimental evidence that network topology can accelerate the spread of beneficial mutations

**DOI:** 10.1101/2021.07.13.452242

**Authors:** Partha Pratim Chakraborty, Louis R. Nemzer, Rees Kassen

**Affiliations:** Department of Biology, University of Ottawa; Ottawa, K1N 6N5, Ontario, Canada; Department of Chemistry and Physics, Halmos College of Arts and Sciences, Nova Southeastern University; Ft. Lauderdale, FL, USA

## Abstract

Whether the spatial arrangement of a population influences adaptive evolution has puzzled evolutionary biologists. Theoretical models make contrasting predictions about the probability a beneficial mutation will become fixed in a population for certain topologies like stars, where leaf populations are connected through a hub. To date, these predictions have not been evaluated under realistic conditions. Here, we test the prediction that topology can change the fixation probability both *in vitro* and *in silico* by tracking the dynamics of a beneficial mutant under positive selection as it spreads through networks of different topologies. Our results provide empirical support that metapopulation topology can increase the likelihood that a beneficial mutation spreads, broadens the conditions under which this phenomenon is thought to occur, and points the way towards using network topology to amplify the effects of weakly favored mutations under directed evolution in industrial applications.

## Main text

How network topology – the pattern of connectivity among subpopulations – impacts adaptive evolution remains poorly understood. Population genetic models tracking the effects of migration and selection on gene frequencies, often under the simplifying assumption of infinite population size, predict little effect of spatial structure on fixation probabilities of beneficial mutations^1,2^. By contrast, models employing evolutionary graph theory (EGT), in which individuals occupy the vertices of a graph and edges represent dispersal routes between neighboring sites, predict that fixation probabilities can change based on how the nodes are connected^3^.

More specifically, EGT predicts that a rooted four-deme network (composed of satellite ‘leaves’ connected by a central ‘hub’) in which one patch supplies more individuals to others than it receives (Fig 1A, left panel) can decrease (suppress) fixation probabilities relative to a well-mixed system (Fig 1A, middle panel). A beneficial mutant is likely to spread through this topology only if it arises in the hub and selection is substantially stronger than migration, consistent with migration-selection models in population genetics. The predictions made by standard population genetic models and EGT are different for networks with connections clustered in a few vertices like a star (Fig 1A, right panel).While population genetic models predict no effect of topology on the rate or probability of spread, EGT predicts fixation probabilities can increase (amplify) compared with a well-mixed system^3,4^ because beneficial mutations arising in a leaf can spread to all other patches via the central hub.

**Figure 1.**
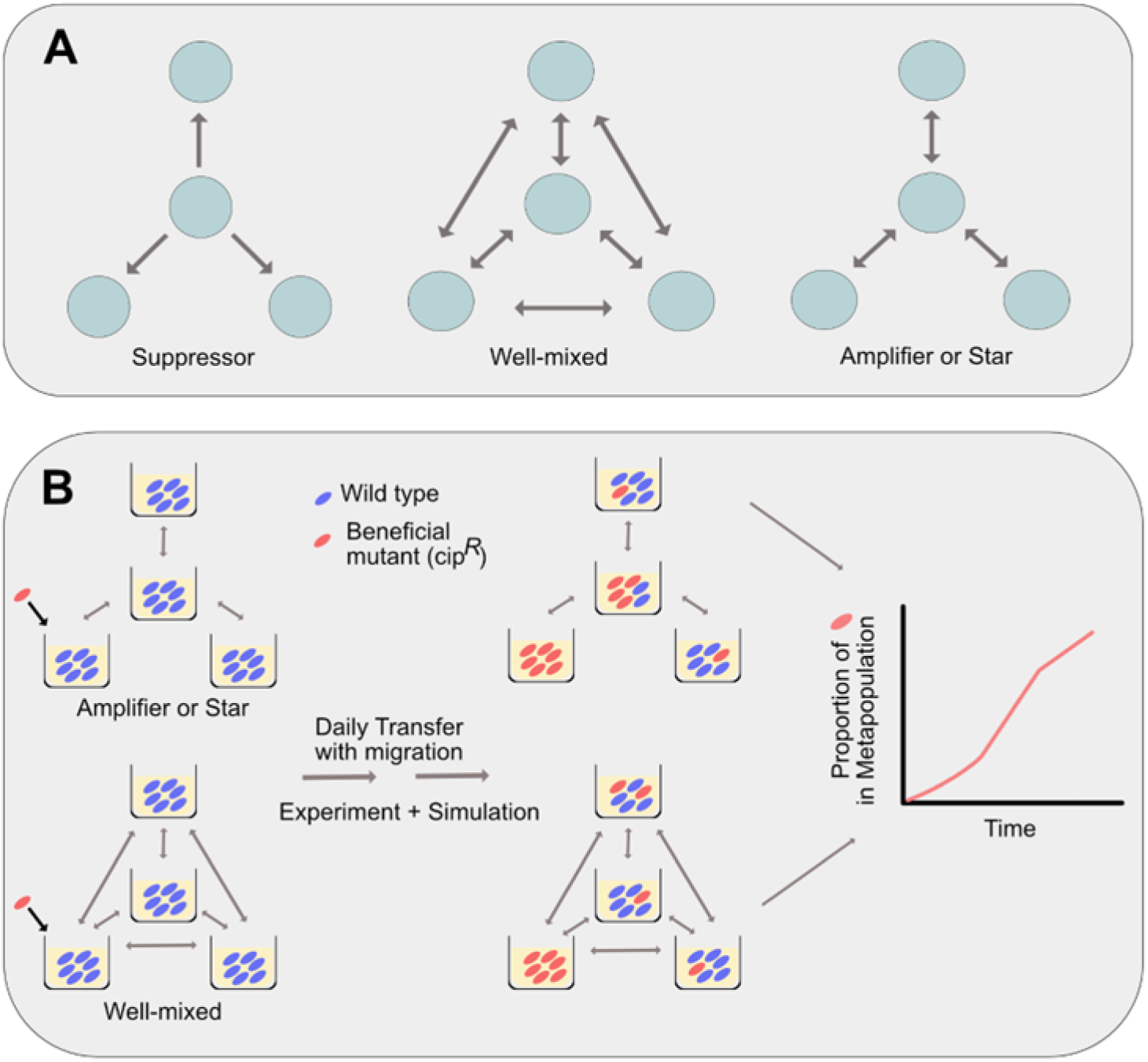
Network Topologies and experimental design: **A.** Three network topologies among four subpopulations. Arrows depict dispersal among subpopulations (green circles). **B.** Experimental schematics.

The EGT approach has inspired a rich theoretical literature exploring the potential for ever more complicated network structures to serve as amplifiers of selection^5^, including claims that certain topologies, like the so-called “superstar”, could asymptotically amplify even very small fitness differences. Later work^6^ tempered these findings by showing that this prediction depends on model details like the ordering of birth and death steps, and that perfect amplification would require unlimited space and time^5^. In any case, the central claims of EGT remain untested by experiment in any biological system, and so its relevance to real-world situations remains uncertain. Moreover, because EGT rests on a stochastic model of evolution in finite populations^7^ where individuals occupy the nodes of the graph, the empirical robustness of its predictions to biologically realistic situations involving nodes composed of subpopulations, variable rates of dispersal, or migration asymmetries^8–12^ remains unclear.

Here, we evaluate the impact of network topology on the probability of fixation of a beneficial mutation directly through experiment. We focus on the spread of antibiotic resistant mutant through a star topology across a range of dispersal rates, as this is the network structure where EGT and standard population genetics make divergent predictions. The well-mixed topology serves as a control. Previous experimental work on the impact of spatial structure on adaptive evolution has considered the impact of population subdivision on the magnitude or extent of adaptive change^13–19^, the emergence and maintenance of diversity^20–23^, and community resilience^24^, rather than the speed or probability of fixation.

Our experiment tracks the spread of an initially rare (1:1000) ciprofloxacin-resistant (cip^R^) mutant of *Pseudomonas aeruginosa* strain 14 (*PA14-gyrA*) invading four-patch metapopulations varying in topology and dispersal rate (Fig 1B). Selection is uniform across all patches, and is imposed by supplementing growth media with subinhibitory concentrations of ciprofloxacin adjusted to provide a ~20% fitness advantage to the resistant mutant. Dispersal occurs during daily serial transfer by first mixing samples from the appropriate subpopulations (see Methods for details) and then diluting the mixture to adjust dispersal rates. Since the theory makes predictions about fixation of a single beneficial mutation, we focus on the first 5 - 6 days (~6.67 generations/day: ~35 - 40 generations) to minimize the opportunity for *de novo* mutations rising to high frequency. We check our experimental results with a new agent-based simulation (see Methods) in which an individual is represented by an agent that competes for finite spaces in a node and can disperse along edges. Together, our results allow us to test directly, both *in vitro* and *in silico*, whether network topology modulates the fixation process that drives adaptive evolution, and if so, how this occurs.

## Results

Our results show that the effect of network topology on the spread of the cip^R^ mutant depends on migration rate (Fig 2). The beneficial mutant spreads faster through a well-mixed than a star topology at migration rates above 10% (panel A: final frequency of cip^R^ in networks with 30% migration: χ2 = 15.348, p<0.0001; 20% migration: χ2 = 9.0148, p = 0.0027) while the rate of spread is statistically indistinguishable at the intermediate migration rates of 10% and 1% (10% migration: χ2 = 0.7082, p = 0.4001; 1% migration: χ2 = 0.2168, p = 0.6415). Below migration rates of 0.01% we see evidence that cip^R^ for modestly faster spread in a star network than in a well-mixed system, consistent with the prediction from EGT that bidirectional star networks can amplify selection. Notably, the amplification effect is transient, being maximal at intermediate time steps (0.01%: relative frequency of cip^R^ to WT on Day 5: χ^2^ = 13.825, p = 0.0002 and 0.001%: frequency of cip^R^ on Day 4: χ^2^ = 5.2577, p = 0.0218) and disappearing on day 5 or 6, depending on the migration rate (see supplementary information), an effect that has not been previously observed in models of EGT.

**Figure 2.**
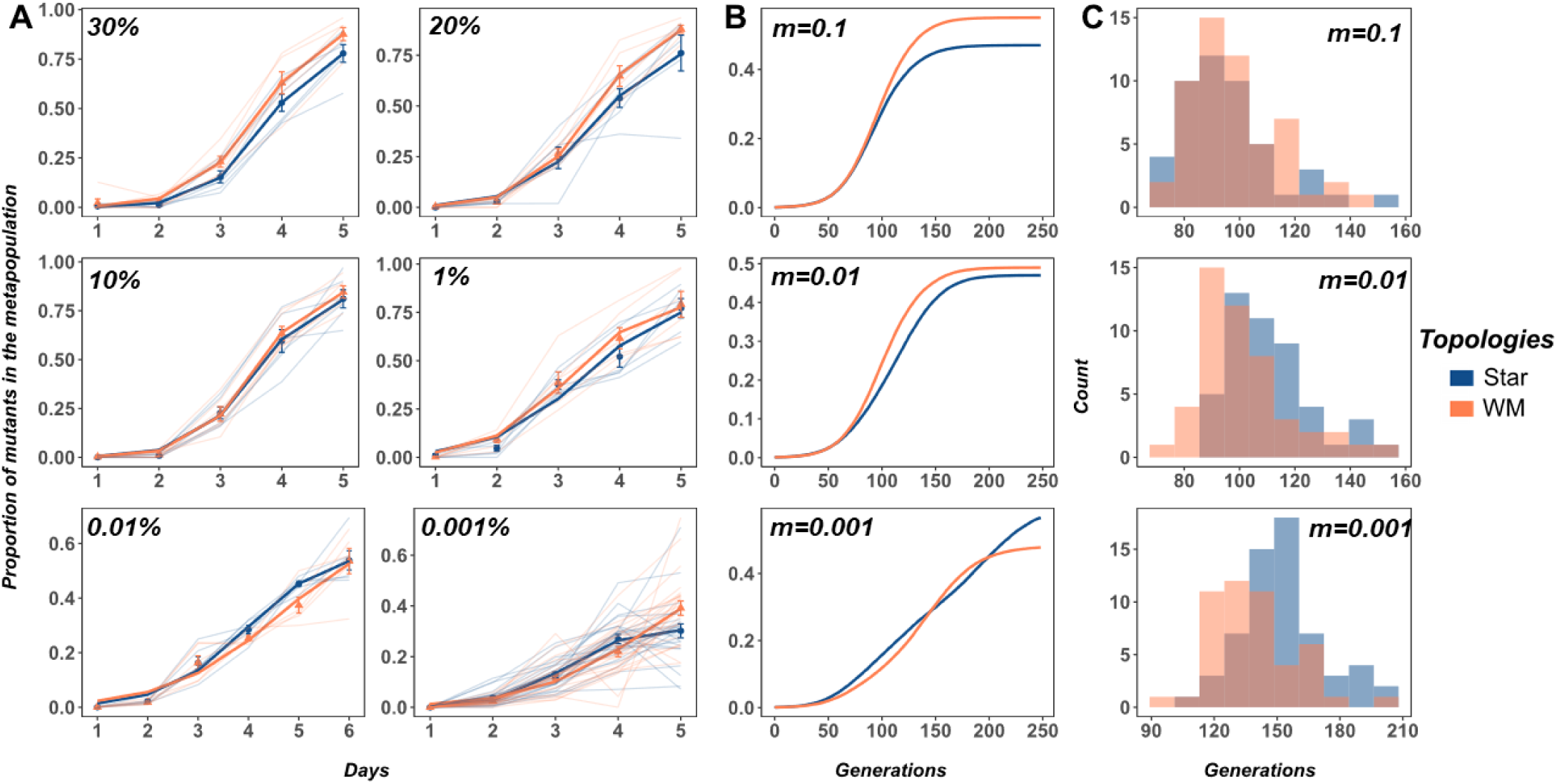
The proportion of cip^R^ mutant in replicate metapopulations propagated by either star (blue) or well-mixed (red) networks with unweighted migration. The bright lines depict the nonlinear least squares (NLS) fit for each metapopulation structure. Panel A shows experimental results; simulation results are shown in Panel B. Migration rates are noted in the inset of each plot. Five days in the experiment is equivalent to approximately 240 generations in the simulation (see methods for details). Panel C are histograms from the simulations showing the minimum generations required for the cip^R^ to reach a frequency of at least 50% in a metapopulation under each combination of network type and migration rate. Each data point represents the mean proportion of all of the replicate metapopulations on that day and error bars represent standard error.

Our results suggest that amplification can occur in star topologies, although its effects appear limited to very low migration rates. An alternative explanation is that amplification in our experiments was due to *de novo* evolution of second-site beneficial mutations in the cip^R^ background. Three lines of evidence argue against this interpretation. First, there should be no inherent evolutionary advantage to any treatment in our experiment because mutation supply rates, being the product of population size and mutation rate, did not differ across treatments. Second, we observed novel colony morphotypes in the more abundant wild-type background only that, when present, had undulate morphologies indicative of biofilm formation. Third, we never recovered mutants more resistant than our focal cip^R^ strain, even after propagating the wild type for ten days under identical conditions (see methods, extended figure 6), suggesting that spontaneous resistant mutations, if present, remained too rare to influence our results. Our observation that amplification of an initially rare beneficial mutant occurred in spite of potential competition from *de novo* mutants in the more abundant competitor class thus makes our results even more compelling.

To confirm these results are not an idiosyncratic feature of our biological system, and to provide additional insight into the mechanisms driving amplification, we simulated the population dynamics of selection in metapopulations under the same topologies and migration rates using an agent-based model. The model tracks competition between wild-type and resistant bacteria for a fixed number of spaces in each patch with dispersal along edges between patches, with fitness being given by the probability of being killed by the antibiotic, and population sizes within each patch being allowed to vary between zero and a fixed carrying capacity. Our model thus allows us to capture the dynamics of slow, but nonequilibrium, migration. Simulation (Fig 2B-C) and experimental results match closely, with the well-mixed topology being faster at spreading the beneficial mutant than the star network at high migration rates. As in our experimental results, transient amplification was seen under low migration rates (Panel B, Fig. 2). Closer inspection reveals amplification is most likely to occur when the expected number of mutant migrants per generation along each edge, at the effective carrying capacity of mutants, is on the order of one. This corresponds to:

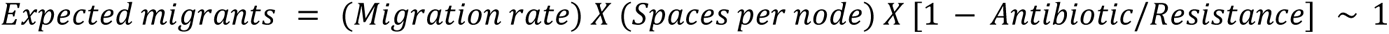

where *Antibiotic* is the antibiotic concentration and *Resistance* represents the reduction in the kill rate of mutants normalized by their growth rate, so that all of them would be killed when (*Antibiotic/Resistance*) ≥ 1. This result suggests that amplification associated with “slow” migration rates is caused by seeding events of mutants into a new subpopulation if and only if the mutants have already successfully colonized a previous subpopulation. Substitution occurs in a more predictable and stepwise fashion under slow, relative to fast, migration rates because beneficial mutants have more time to rise to high frequency in the subpopulation they previously colonized, and so are less likely to be lost due to drift when colonizing a new subpopulation. Moreover, a star network that concentrates incoming migrants into the hub will alleviate genetic drift more than a well-mixed network, and should, in principle, act as a stronger amplifier of selection. By contrast, when migration introduces beneficial mutants at a rate faster than they are lost due to genetic drift, a better connected well-mixed network spreads beneficial mutants faster than a star network where leaves are only connected via the hub.

Recent theoretical work^9,25^ treating nodes as subpopulations rather than individuals, as in our experiments, shows that migration asymmetry between leaves and hub can potentially amplify fixation probabilities relative to the well-mixed case. Specifically, star networks with net outward or inward migration are predicted to be suppressors or amplifiers of selection, respectively, while those with no asymmetry (balanced) migration should have no advantage over a well-mixed network in fixing a beneficial mutant^9^. Our first experiment (Figure 2) adjusted the migration rate, *m*, to ensure all patches receive the same number of mutants. Using the same experimental setup, we can test these predictions by manipulating the relative amount of migration between the hub and the three leaves of the star network in both experiments and simulation.

Our results are consistent with the predictions of theory. At high migration rates (Fig 3A), the rate at which the cip^R^ mutant spreads is never statistically significantly higher than that of the well-mixed topology under both forms of asymmetric migration (middle and bottom panels; χ^2^ = 1.1888, p = 0.2756 (OUT>IN) and χ^2^ = 0.4124, p = 0.5208 (IN>OUT), respectively; see supplementary information), and is substantially slower when migration rates were balanced among the nodes (top panel) (IN=OUT: relative frequency of cip^R^ on Day 9: χ^2^ = 12.234, p = 0.0005). At low migration rates (Fig 3B), however, the dynamics of cip^R^ spread is indistinguishable (see supplementary information) from that of the well-mixed case for both balanced migration (top panel, χ^2^ = 0.1848, p = 0.6673) and net outward migration (middle panel, χ^2^ = 0.5233, p = 0.4694), as expected from theory. When inward migration exceeds outward migration (bottom panel), however, the cip^R^ mutants gain a significant advantage in the latter stages of the experiment (IN>OUT: relative frequency of cip^R^ on Day 9: χ^2^ = 5.2917, p = 0.0214). The results of our simulations are shown in the panels in Fig 3C and match our experimental results closely.

**Figure 3.**
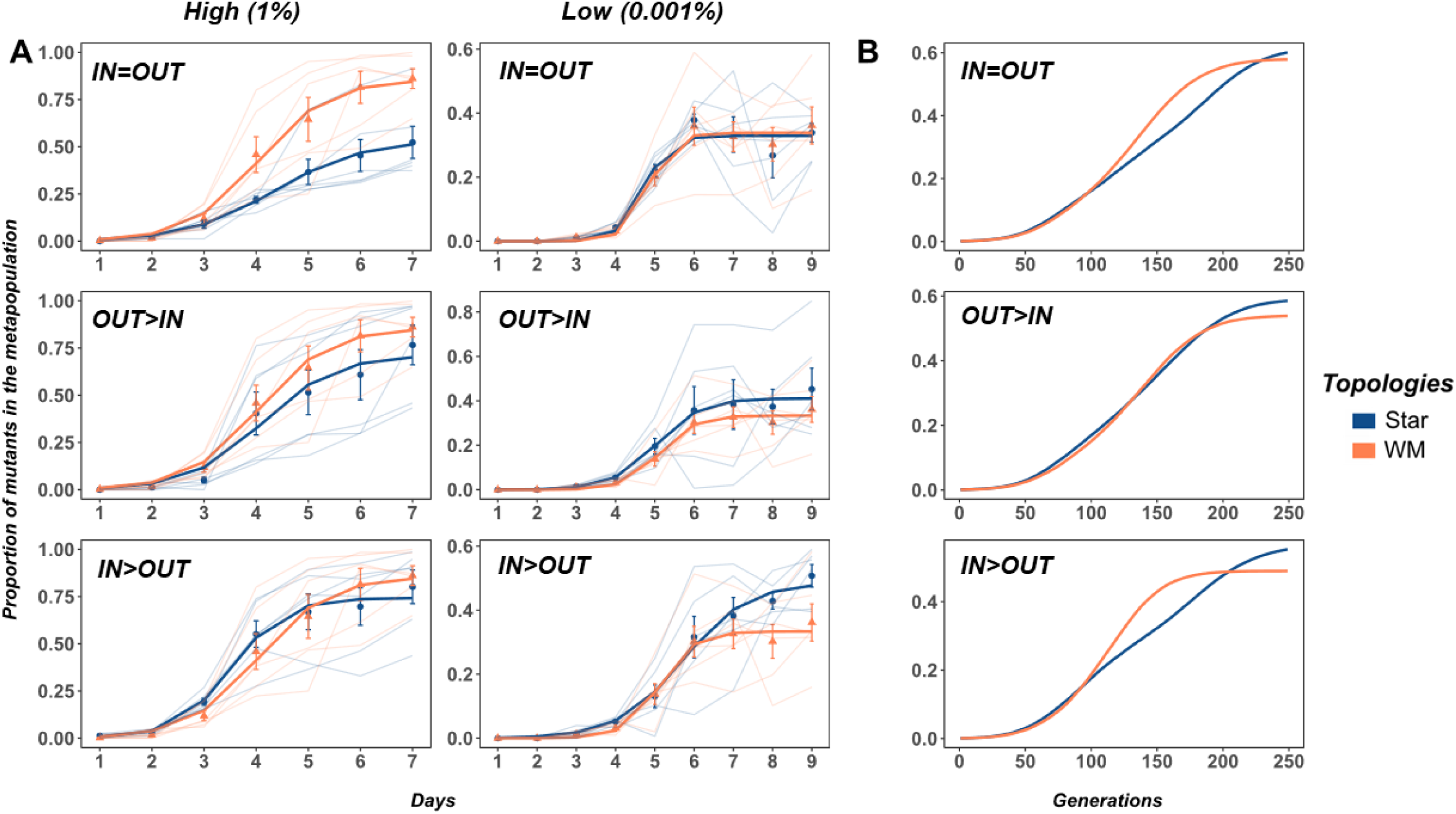
The proportion of cip^R^ mutant in replicate metapopulations propagated on either star (blue) or well-mixed (red) networks with weighted migration. Bright lines are the nonlinear least squares (NLS) fit to the two network treatments. Panel A shows results from the experiments under high and low migration rates, whereas Panel B shows the results of the agent based model only under the low migration rate. The respective dispersal asymmetries are provided in the inset of each plot. Each data point represents the mean proportion of all of the replicate metapopulations on that day and error bars represent standard error.

Whether or not amplification occurs should depend on the balance between two dynamic processes - the rate of fixation within a patch and the rate of dispersal to new patches. When migration rates are fast, an initially rare beneficial mutant cannot reach sufficiently high frequency in its native patch to guarantee dispersal to other patches. Consequently, if it does manage to get dispersed to a novel patch, the beneficial mutant is initially so rare that it is likely to be lost due to drift. Under slow migration, however, selection increases the frequency of a beneficial mutant faster in its native patch than it is dispersed to novel patches, ensuring that it can be repeatedly dispersed to novel patches and so reducing the likelihood of drift loss upon arrival. Amplifying topologies act in a similar way when dispersal is slow, by allowing beneficial mutants to first fix in the patch where they were introduced, and then funneling them through a central hub, so the likelihood of drift loss before other leaves are seeded is reduced. Under well-mixed conditions, migration overwhelms selection such that the constant influx of lower fitness migrants from other patches means the beneficial mutant cannot accumulate to sufficiently high frequency in its focal patch before it is dispersed to other patches, where it is rare and likely to be lost due to drift.

We evaluated this interpretation by examining the dynamics of the cip^R^ mutant as it spreads among subpopulations in our experiment. Fixation is expected to occur first in the leaf in which the beneficial mutant was initially inoculated followed by, in an amplifying star network, accumulation in the hub and then spread to other leaves of the network. In a well-mixed metapopulation, however, the spread of the cip^R^ mutant should occur into both hub and leaf subpopulations at the same time. Indeed, when we examine the dynamics of the cip^R^ mutant in star and well-mixed metapopulations at a low migration rate (0.001%) where amplification is seen in the former but not the latter, we see the expected patterns (Fig 4). The cip^R^ mutant first fixes in the leaf where it was initially inoculated for both of the networks, as expected. Although there is substantial variation among replicates, our results show that the cip^R^ mutant spreads differently among the remaining three subpopulations in the two kinds of network: in the star network there is a clear tendency for the mutant to spread from the initial subpopulation to the hub (P2) first, whereas in the well-mixed network the mutant is equally likely to spread to the hub as any additional subpopulation (Fig 4A). These experimental results are mirrored closely by those of the simulation (Fig 4B).

**Figure 4.**
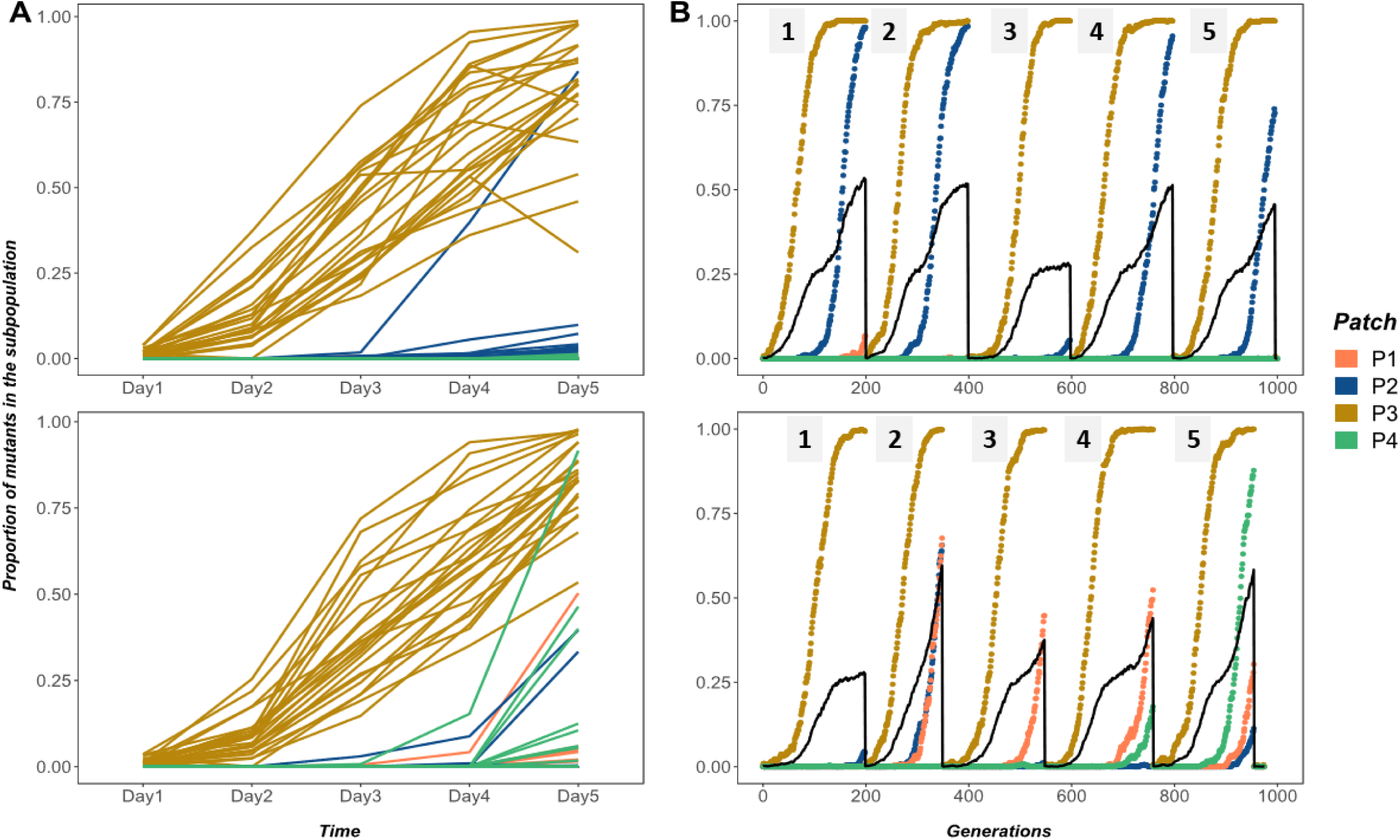
The proportion of the cip^R^ mutants in the constituent subpopulations of each replicate metapopulation propagated as either star (both panel A and B, first row) or well-mixed (both panel A and B, second row) networks with unweighted migration of 0.001% (~100 individuals). Panel A is the data from the experiment and Panel B is the data from the agent based model. Subpopulation nomenclature: P3 = node of introduction of the cip^R^ mutant, P2 = hub and P1 and P4 = rest of the peripheral leaves. In panel A, cip^R^ fixation dynamics in four subpopulations of each of the 24 replicate metapopulations under each network are shown (top = star, bottom = well-mixed). The black solid lines in panel B are the overall proportion of cip^R^ mutants in a metapopulation. In panel B, five replicate instantiations (each run for 200 generations) of the simulation are shown for each network (top = star, bottom = well-mixed).

We see similar dynamics of spread among subpopulations in our experiments examining migration asymmetry. In well-mixed systems and those star networks where amplification was not observed (high migration rates) the cip^R^ mutant rapidly spreads into both hub and leaves after near fixation in the patch of introduction (Fig 5A). In contrast, the cip^R^ mutant spreads to the hub first in the most strongly amplifying star topology (IN>OUT), consistent with the idea that beneficial mutants are more likely to avoid stochastic loss due to drift by being concentrated in the hub (Fig 5B).

**Figure 5.**
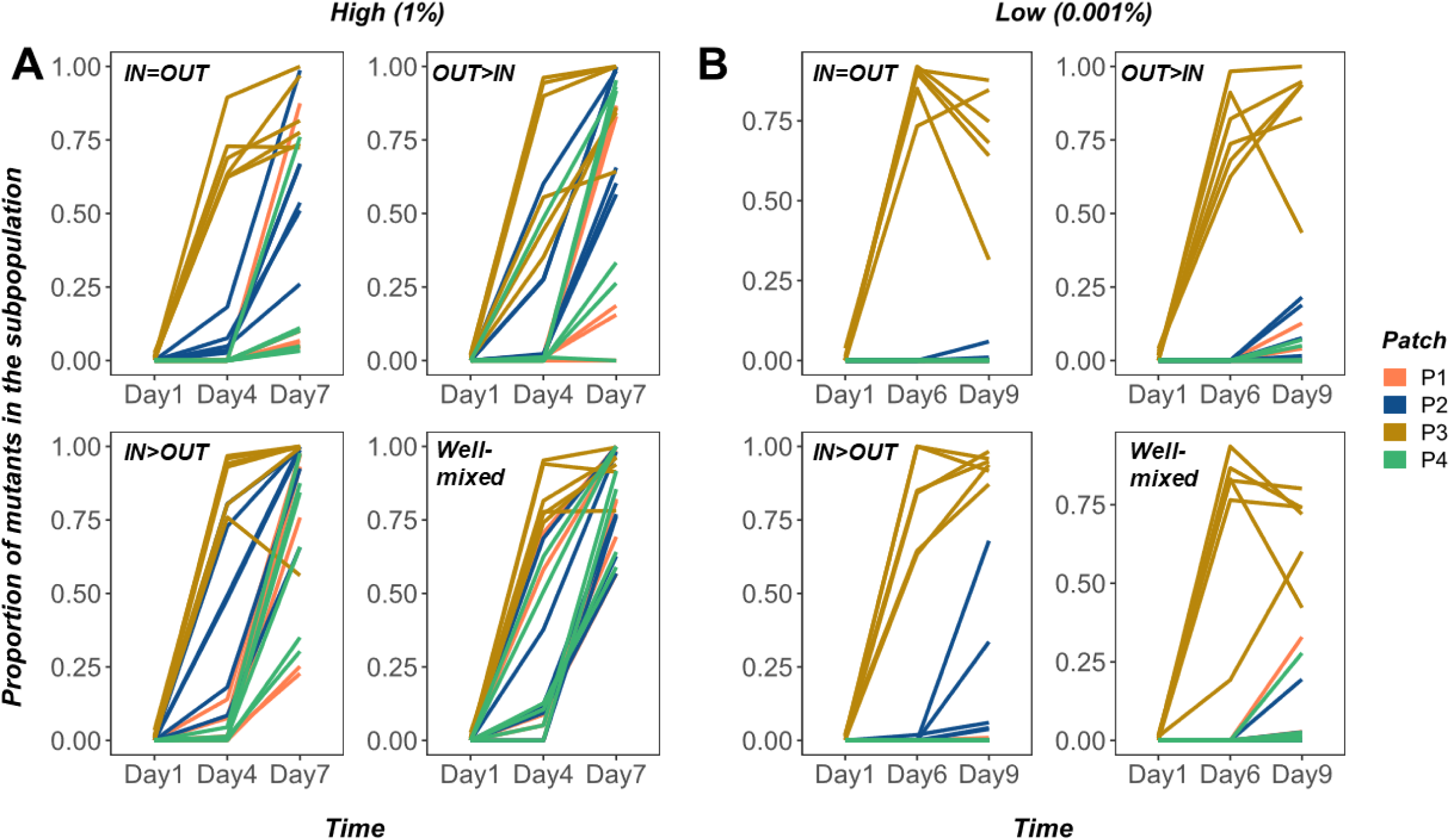
The proportion of the cip^R^ mutants in the constituent subpopulations of each metapopulation propagated by either asymmetric star or well-mixed networks under high (panel A: 1% or 10^5^ individuals) and low (panel B: 0.001% or 10^2^ individuals) weighted migration. Subpopulation nomenclature: P3 = node of introduction of the cip^R^ mutant, P2 = hub and P1 and P4 = rest of the peripheral leaves. In both panels A and B, cip^R^ fixation dynamics in four subpopulations of each of the six replicate metapopulations using each of the three asymmetric star networks and the well-mixed network are shown (see plot insets for details) under high and low migration rates, respectively.

If avoiding “drift loss” under low migration rate is indeed the mechanism for amplification, then imposing more severe drift should increase the magnitude and duration of the observed amplification. We tested this hypothesis by enforcing a stricter bottleneck, and thus stronger drift, during daily transfers and tracking the spread of resistant mutants in a star metapopulation with in-weighted migration (IN>OUT) and a well-mixed metapopulation at a low migration rate (~1000 individuals). The results are consistent with our prediction (Fig 6): for intermediate time-points (Day 5 and 6), there was a significantly higher proportion of mutants (Day5: χ^2^=17.255, p = 3.269e-05, Day6: χ^2^=16.17, p = 5.79e-05) in the star-like metapopulations compared to the wild-type. In other words, we observed an amplification with a higher magnitude and longer duration (stable for ~30 generations, which was nearly the entire length of our previous experiments). This result is consistent with our hypothesis that the likelihood of drift loss is lower in star-like metapopulations when the migration rate between subpopulations are low and asymmetric migration concentrates mutants through the hub. Our results thus lend strong support to the idea that a reduction in the probability of drift loss is responsible for the amplification effect in star-like topologies. Moreover, this result emphasizes the need for future theoretical and experimental work to focus on fine-tuning evolutionary forces such as stochastic drift, selection and migration to determine the magnitude of amplification in different topologies.

**Figure 6.**
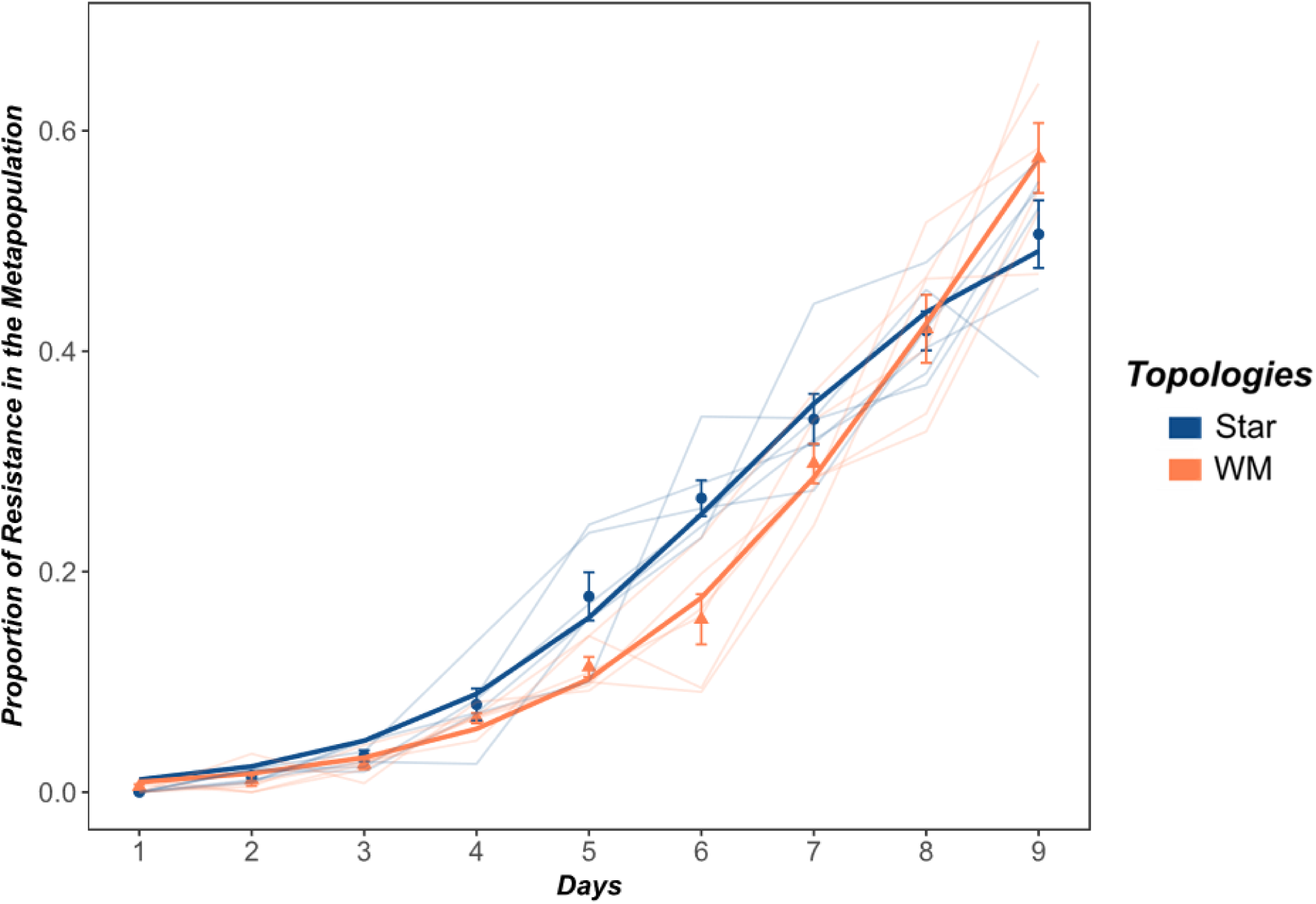
The proportion of cip^R^ mutant in replicate metapopulations propagated on either an inward star (blue) or well-mixed (red) networks with low population size (10^5^ CFU/mL) and low migration rate (10^3^ CFU/mL). Bright lines are the nonlinear least squares (NLS) fit to the two network treatments. Each data point represents the mean proportion of all of the replicate metapopulations on that day and error bars represent standard error.

## Discussion

We have shown experimentally and through simulation that metapopulation structure can impact the dynamics of adaptive substitution. Star metapopulations, in which leaf subpopulations exchange migrants through a central hub, can act as amplifiers of selection leading to faster rates of spread than a comparable well-mixed population where all subpopulations share migrants equally. Amplification is most pronounced when selection is strong relative to migration, a scenario that reduces the probability that a rare beneficial mutant in a newly colonized patch will be lost due to drift, and when the topology of dispersal concentrates the beneficial mutant in a central hub (‘inward’ > ‘outward’ migration). In other words, amplification occurs because rare mutants are less likely to be lost, not because the strength of selection itself increases.

This result is remarkable because it was not anticipated by standard theory in population genetics, in which population structure usually has little effect on the probability of fixation for beneficial mutants. This conclusion likely derives from the tradition in population genetics of considering allele frequency changes in the limit of infinite populations and high migration rates. Our results, by contrast, show how stochastic effects associated with finite population sizes can alter the dynamics of adaptive substitution in ways that are consistent with predictions from evolutionary graph theory where individuals are assigned to nodes of a graph. Importantly, our results show that amplification can occur under a broader and arguably more realistic set of conditions where populations, not individuals, occupy the nodes. Our work emphasizes the previously overlooked importance of migration rate and serves as a first step towards bridging these two approaches, with infinite populations on the one hand and finite populations focused on the dynamics of individuals on the other.

More generally, it will be useful to expand the analytical framework of EGT to include more biological realism and to articulate more precisely the range of conditions under which amplification can occur. It should be possible, for example, to use network topology to amplify the selection of even a slightly favored mutation for the purpose of experimentation or the directed evolution of desired traits in industrial applications. A more comprehensive theory of evolution on structured landscapes will also be important in other aspects of biology, including the spread of invasive species, pathogens, and the resistance factors they possess.

## Methods

### The SANCTUM model

**Supplementary Figure 1:**
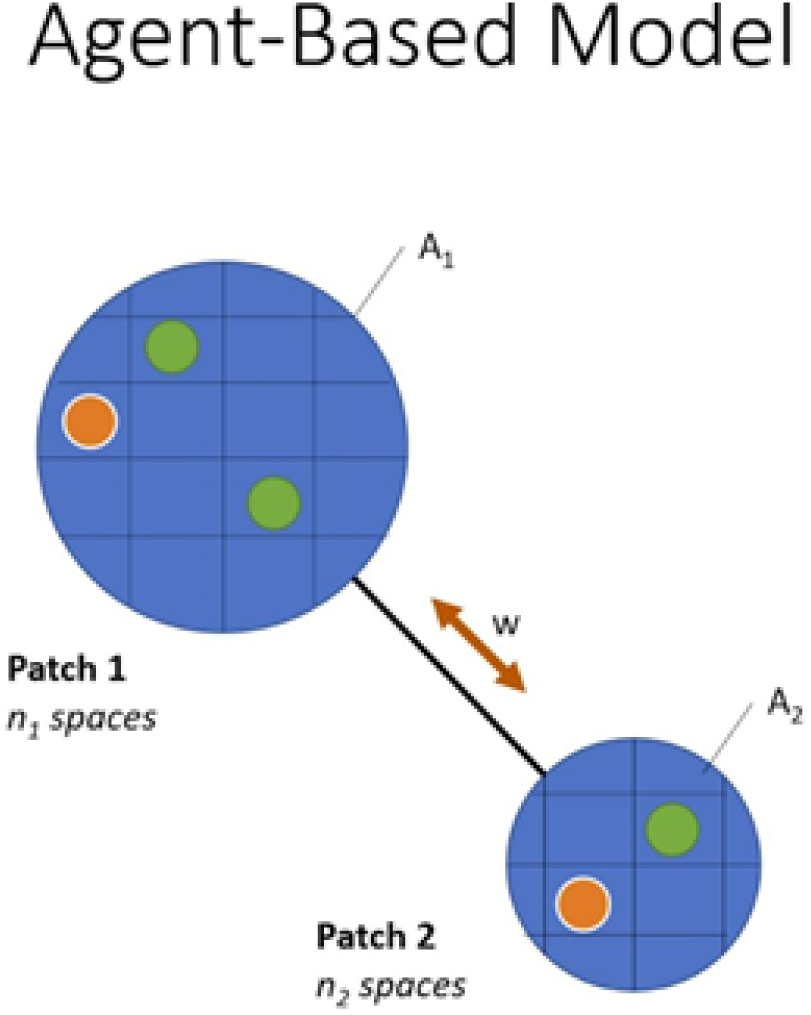
Schematic depiction of the algorithm used in the SANCTUM model. See text for details.

Each node (Ai) has ni spaces that can each be empty, occupied by a wild-type, or occupied by a mutant. For these experiments, all the n values are set to 1600. For each generation of the simulation, there are three phases: (1) Death, (2) Birth, and (3) Migration. *Death:* An agent is removed during the death step with a probability that depends on the antibiotic concentration divided by its individual resistance. *Birth:* Each agent has a chance of reproducing an identical agent in an empty space of the same node during the birth step. Similar to the Lotka-Volterra model of population growth, the probability of reproduction increases with the number of empty spaces in the node. *Migration*: There is a probability of migration that varies based on the experimental condition. If an agent is selected to migrate, it randomly moves to an adjacent connected node with a probability proportional to the weight of that edge (w). These definitions are interpreted in the model as follows:

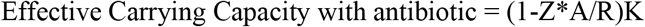

where Z is first-order kill rate of the mutants by the antibiotic, A is the antibiotic concentration, R is growth rate of mutants, and K is the total number of spaces per node (the unmodified carrying capacity). Using the effective carrying capacity of mutants limited by the antibiotic:

Expected migrants per step per edge = (Migration rate) x (Effective Carrying Capacity)

= (Migration rate) x [(1-Z*A/R)K]

= (Migration rate) x (Spaces per node) x [1-Antibiotic/Normalized Resistance]

Here, R/Z represents the Normalized Resistance, which reflects the balance between the death and growth rates. This value also sets the threshold for A, above which all the mutants would be eliminated. The initial system is randomly seeded with one thousand agents across all nodes, and one of these is selected to be the mutant (not in the hub). For each simulated condition, the metapopulation fraction is averaged over 100 instantiations. For runs that ultimately fix, the number of generations until the mutants are the majority of agents is also recorded.

### Microbial strains and conditions

For all experiments, clonal populations of *Pseudomonas aeruginosa* strain 14:gyrA (*PA14:gyrA*) and *PA14: lacZ*, isogenic to PA14 except with a point mutation in the *gyrA* gene and an insertion in the *lacZ* gene respectively, were used. Colonies possessing the *lacZ* insertion appear blue when cultured on agar plates supplemented with 40 mg/L of 5-bromo-4-chloro-3-indolyl-beta-D-galactopyranoside (X-Gal), and are visually distinct from the PA14: *gyrA* white colouration. The neutrality of the *lacZ* marker was confirmed in our experimental environments by measuring the fitness of the marked strain relative to the unmarked strain. Populations were cultured in 24-well plates with 1.5 mL of media in each well, in an orbital shaker (150 RPM) at 37°C. The culture media consisted of Luria Bertani broth (LB: bacto-tryptone 10 g/L, yeast extract 5 g/L, NaCl 10 g/L) supplemented with 20 ng/mL of the fluoroquinolone antibiotic, ciprofloxacin. This concentration of ciprofloxacin confers ~ 20% selective advantage to *PA14:gyrA* relative to *PA14:lacZ* (extended figure 5). All strains and evolving populations were frozen at −80°C in 20% (v/v) glycerol.

### Evolution experiment

A single metapopulation consisted of four subpopulations, one subpopulation being located on each of four different 24-well plates. Plate 2 was always assigned as the hub, and plates 1, 3, and 4 were treated as the leaves. This design allows us to track up to 24 replicate populations using just four multi-well plates. The experiment was initiated by inoculating each subpopulation with ~10^7^ colony forming units (CFU) per ml of PA14:*lacZ* descended from a single colony picked from an agar plate and grown overnight in liquid LB at 37°C with vigorous shaking (150 RPM). The cip^R^ mutant, derived from frozen cultures in the same way, was introduced simultaneously into one subpopulation (plate 3) at a density ~10^4^ PA14:*gyrA* cells producing an initial ratio of resistant to wild-type cells of ~1:1000 in this subpopulation. Metapopulations were transferred daily following dispersal among subpopulations (see below) by taking an aliquot corresponding to ~10^7^ CFUs per mL and inoculating into fresh medium. The population density in each subpopulation reached ~10^9^ CFUs, so this transfer regime corresponds to ~6.67 daily generations of growth.

We constructed distinct network topologies by mixing subpopulations prior to serial transfer following the schematic shown in Extended Fig 2. Briefly, well-mixed networks were created by combining equal volume aliquots from all subpopulations into a common dispersal pool, diluting this mixture to the appropriate density to achieve the desired migration rate, and then mixing the dispersal pool with aliquots from each subpopulation (so-called ‘self-inoculation’) before transfer. Star networks, which involve bidirectional dispersal between the hub and leaves, were constructed in a similar way to the well-mixed situation only now the dispersal pool consisted of aliquots from just the leaves and aliquots from the hub (plate 2) were mixed with ‘self-inoculation’ samples from each leaf prior to serial transfer. Further details on how each network topology and migration rate were achieved are provided below.

**Supplementary figure 2:**
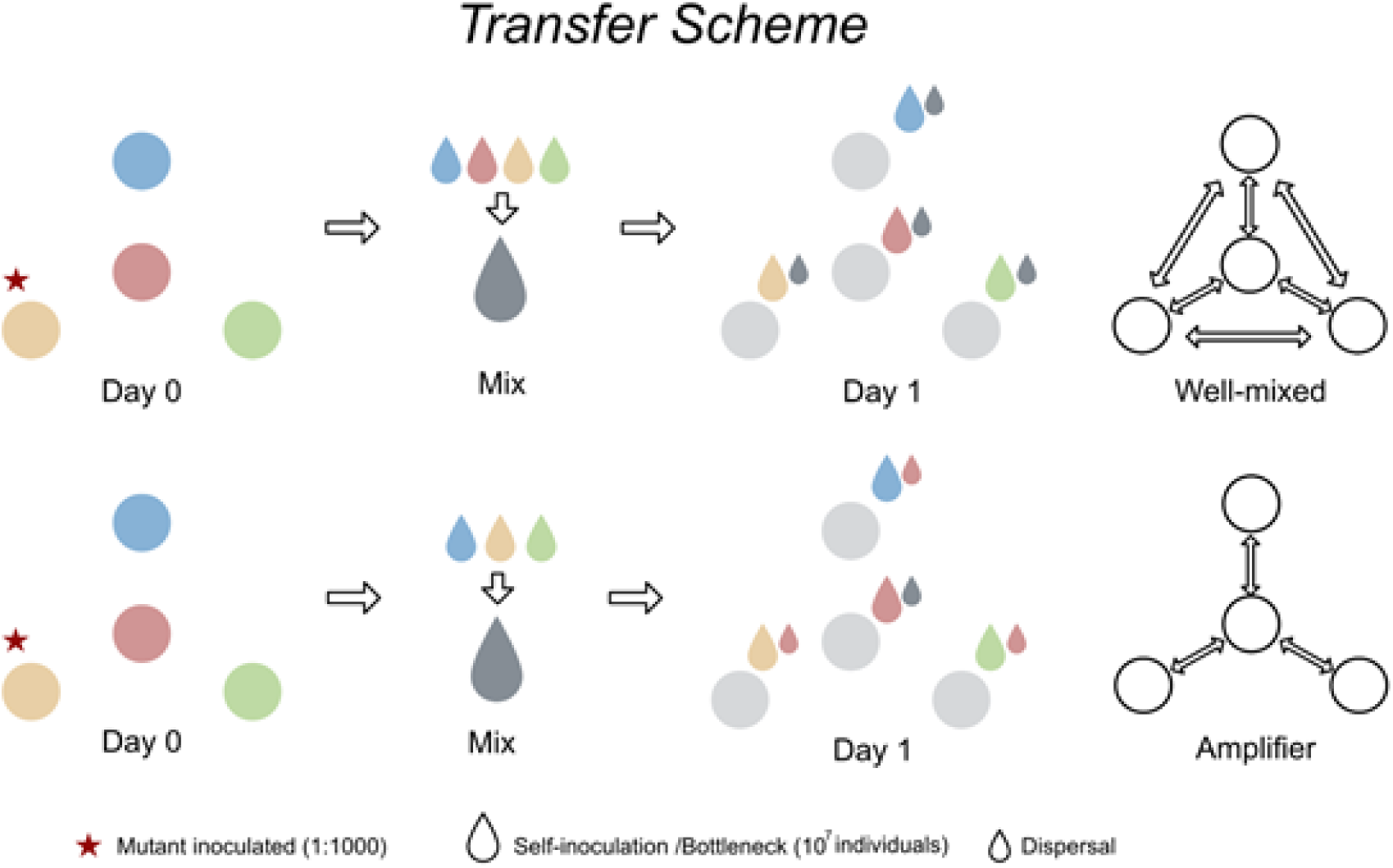
Transfer scheme to experimentally create star and well-mixed network structures. The red star indicates the patch (P3) where the mutant (PA14-*gyrA*) was inoculated 1:1000 ratio to the wild type (PA14-*LacZ*). Big droplets of the four colors indicate self-inoculation from respective patches whereas big and small gray droplets indicate dispersal mix and dispersal volume, respectively.

### Tracking the spread of resistance

We tracked the spread of the cipR mutant (PA14: *gyrA*) relative to the wild type (PA14: *lacZ*) by plating samples from each subpopulation as well as a mixture of the entire metapopulation on LB agar plates supplemented with X-gal allowing us to use blue-white screening to track the relative frequency of each type over time.

### Statistical analyses

All statistical analyses were conducted using R statistical software^26^. We used two complementary approaches to analyze our experimental data.

The first models the spread of resistance (see Fig. 2 and Fig. 3) as a three-parameter logistic growth model using non-linear least squares with a fixed N0 (NLS)^27^. This model, as with comparable approaches focused on population growth in resource limited environments, allows us to estimate the rate at which resistance spreads (equivalent to rmax in logistic growth) and the final frequency of cip^R^ mutants at the end of the experiment (equivalent to the carrying capacity, K, from logistic growth models) in each replicate metapopulation. Contrasts of maximal growth rates between treatments (star or well-mixed) were performed using a linear model (lm function from base R). Comparable contrasts for maximum proportion of resistant mutants fixed on the final day of the experiment used a generalized linear mixed model using methods as described below.

The second approach modeled the change in proportion for the cip^R^ mutants directly using a generalized linear mixed model (GLMM) with quasi-binomial error distribution (and logit link function), using the glmmPQL function from the MASS package in R^28^. We focus on the main effects of time and network structure (star vs. well-mixed) and their interaction at each migration rate treatment. Logistical constraints prevented us from conducting experiments that manipulate both network structure and migration rate simultaneously, so we elected to run separate experiments at each migration rate to focus on the effect of contrasting network structures, as this is the focus of EGT. Our model treats ‘network’ as a fixed effect and ‘replicate’ as a random effect, while accounting for repeated measures through time. This approach produces estimates of the pairwise difference between the slopes (vs time) for the network treatment (for example, - Time:Network star - Time:Network well-mixed) that were further analyzed using the EMTRENDS function from the EMMEANS package (analogous to a Tukey *post hoc* test)^29^. These contrasts allow us to determine the magnitude and direction of difference between the star and well-mixed networks for the whole experiment.

The approaches above, which focus on estimating best-fit main effects and interactions, are useful for helping to visualize the dynamics of spread across many instantiations of an inherently noisy process. We additionally focus attention on contrasts between the fraction of cip^R^ mutants between star and well-mixed treatments at specific days when: (1) the fitted logistic model for the star was higher than that of the well-mixed over the course of the complete experiment; or, (2) when the fitted models reveal a transient “crossover” event at intermediate time steps. We used a GLMM as described above to contrast the fraction of cip^R^ mutants in star vs well-mixed networks at a particular day, treating replicate as a random factor. The analysis of variance of the GLMMs were performed with the ANOVA function from the CAR package.

Full_model <-glmmPQL(Proportion~Time*Treatment, random = 1|Replicate, family = quasibinomial, data)

em1< - emtrends(Full_model, pairwise ~ Treatment, var = ‘Time’)

em1$contrasts

model_K < - glmmPQL(Proportion ~ Treatment, random= ~1|Rep, family = quasibinomial, data) #takes data from final day of the experiment/ change the data to get any particular day

model_R< - lm(R~Treatment, data)

### Detailed methods used to construct network topologies

#### Unweighted migration

**Supplementary Figure 3:**
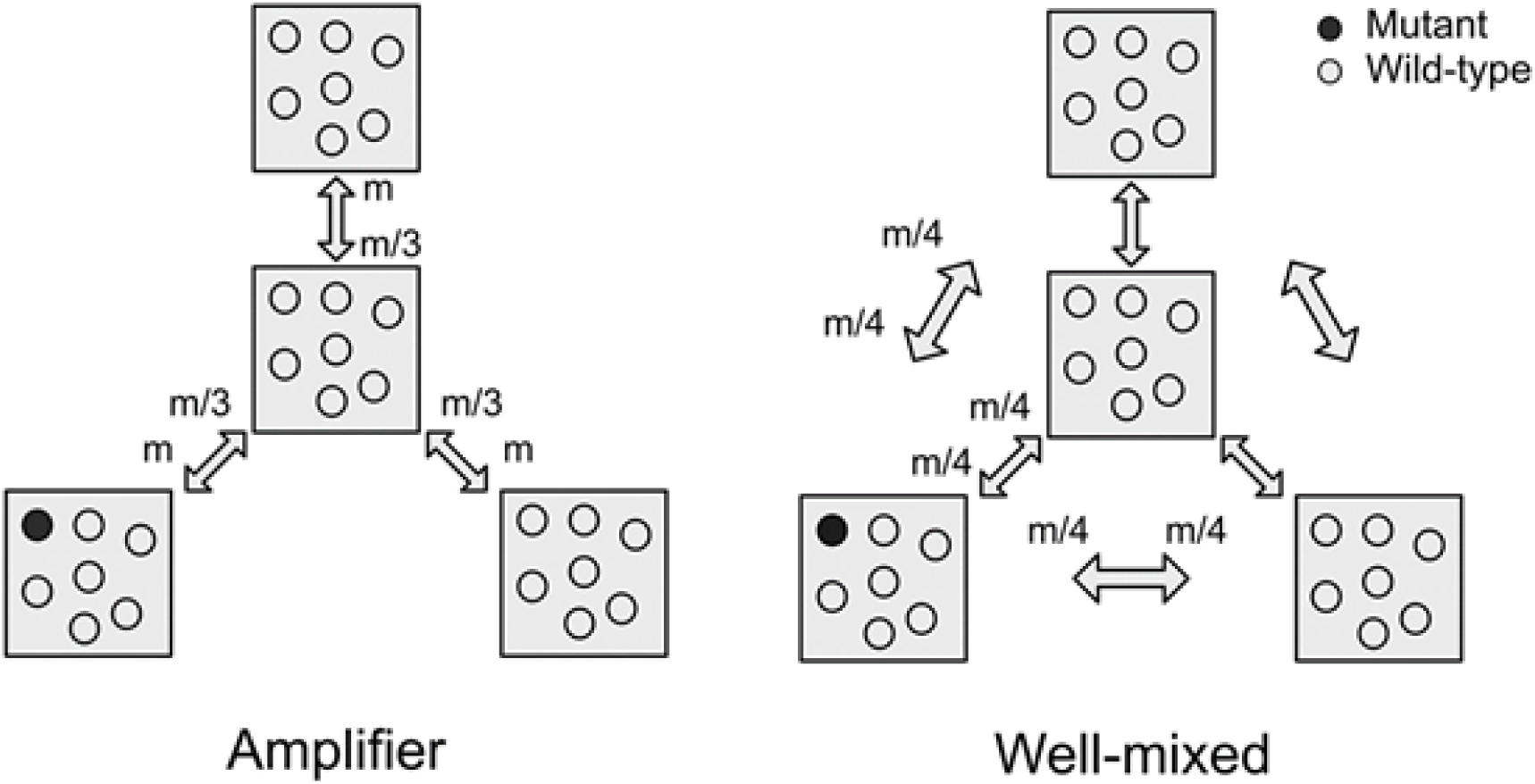
Schematics depicting the number of transferred migrants per edge connection in the cases of unweighted migration regime for star and well-mixed networks. The value of *m* shows the total number of migrants for each migration treatment, and the fractions are calculated as the contributions from each patch. Filled circles (black) indicate the mutants (PA14-*gyrA*) while initializing the experiment and open circles (clear) are the predominant wild type (PA14-*LacZ*).

##### Well-mixed

35 μL of subpopulations 1, 2, 3, and 4 were mixed together, and 20 μL from this resulting pool of migrants (MIX) was serially diluted in fresh media supplemented with 20 ng/mL Ciprofloxacin to achieve ~10^8^, ~10^7^, ~10^5^ and ~10^4^ CFU/mL. Then, 15 μL of the diluted MIX was added to 1.5 mL of fresh media along with 15 μL of the previous day’s culture to achieve desired migration levels of ~10^6^, ~10^5^, ~10^3^ and ~10^2^ CFU/mL. For 20% and 30% migration, 30 μL and 45 μL of the diluted MIX were added with the culture from the previous day.

##### Amplifier

35 μL of subpopulations 1, 3, and 4 were mixed together, and 20 μL from this resulting pool of migrants (MIX) was serially diluted in fresh media and 20 ng/mL antibiotic to reach ~10^8^, ~10^7^, ~10^5^ and ~10^4^ CFU/mL. Also, 20 μL of subpopulation 2 (HUB) was serially diluted in fresh media and 20 ng/mL Ciprofloxacin to reach ~10^8^, ~10^7^, ~10^5^ and ~10^4^ CFU/mL. Then, 15 μL of the diluted MIX was added to 1.5 mL fresh media along with 15 μL of the previous day’s subpopulation 2 culture to achieve the desired migration levels of ~10^6^, ~10^5^, ~10^3^ and ~10^2^ CFU/mL. Also, 15 μL of the diluted HUB was added to 1.5 mL fresh media along with the previous day’s culture to achieve desired migration levels of ~10^6^, ~10^5^, ~10^3^ and ~10^2^ CFU/mL in subpopulations 1, 3, and 4. For 20% and 30% migration, 30 μL and 45 μL of the diluted MIX were added after the bottleneck (“self-inoculation”).

#### Asymmetric migration

**Supplementary Figure 4.**
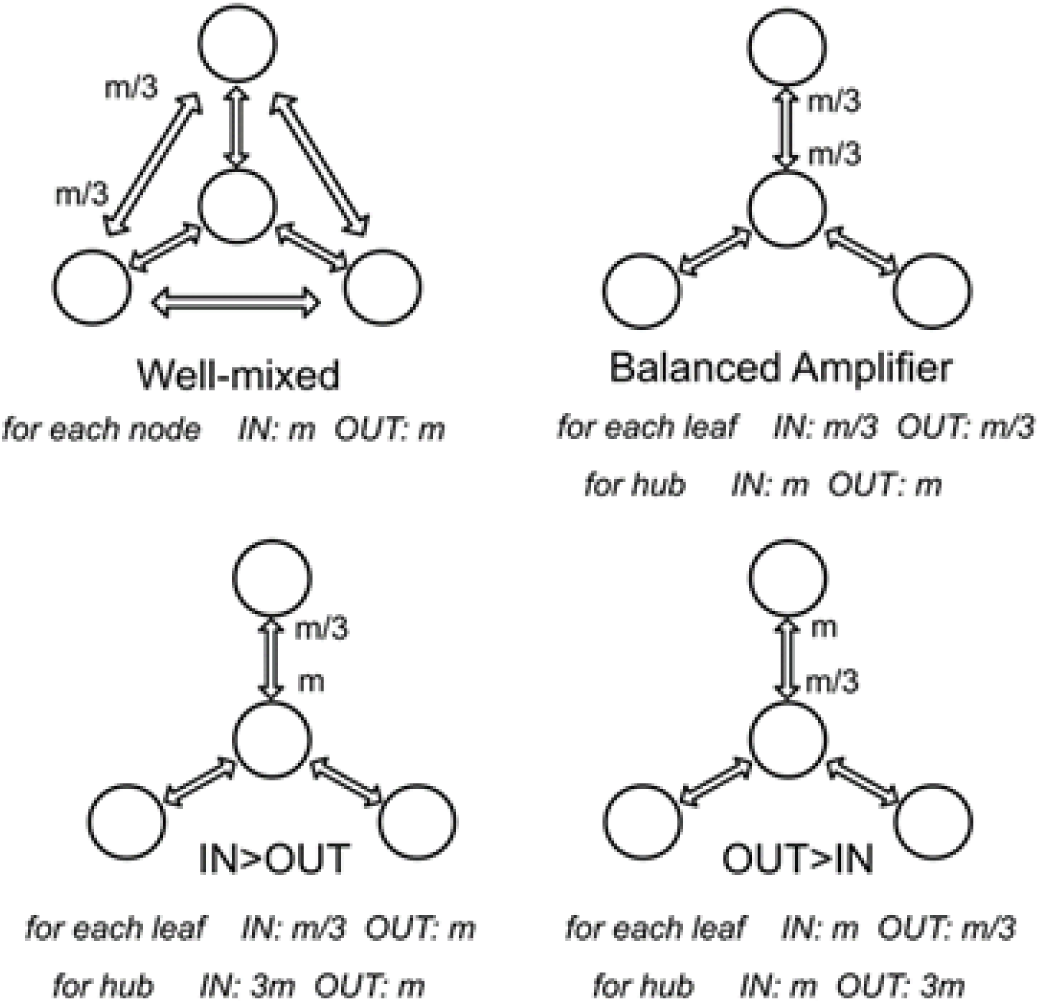
Schematics depicting the transferred migrants per edge connection in the case of the weighted migration regimes for three asymmetric star networks and the well-mixed network. Each double-sided arrow indicates the number of migrants received and contributed by each patch.

##### Well-Mixed

20 μL of subpopulations 1, 2, 3, and 4 (MIX) were mixed in 120 μL of fresh culture media with 20 ng/mL Ciprofloxacin and serially diluted to reach ~4 x 10^7^ or ~4 x 10^4^ CFU/mL. Then, 15 μL of the diluted MIX was added to 1.5 mL fresh media along with 15 μL of the previous day’s culture. This resulted in the transfer of 4 x 10^5^ for the high migration rate experiments, or 4 x 10^2^ CFU/mL for the low migration rate.

##### Balanced star (IN = OUT)

20 μL of subpopulations 1, 3, and 4 (MIX) were mixed in 140 μL of fresh culture media with 20 ng/mL Ciprofloxacin and serially diluted to reach ~3 x 10^7^ or ~3 x 10^4^ CFU/mL. Also, 20 μL of subpopulation 2 (HUB) was mixed in 180 μL of fresh culture media with 20 ng/mL Ciprofloxacin and serially diluted to reach ~10^7^ or ~10^4^ CFU/mL. This resulted in the transfer of ~3 x 10^5^ and ~3 x 10^2^ CFU/mL to the hub, and ~10^5^ and ~10^2^ CFU/mL to the peripheral leaves, for the high and low migration rates, respectively.

##### OUT>IN regime

20 μL of subpopulations 1, 3, and 4 (MIX) were mixed in 140 μL of fresh culture media with 20 ng/mL Ciprofloxacin and serially diluted to reach ~3 x 10^7^ or ~3 x 10^4^ CFU/mL. Also, 60 μL of subpopulation 2 (HUB) were mixed in 140 μL in fresh culture media with 20 ng/mL Ciprofloxacin and serially diluted to reach ~3 x 10^7^ or ~3 x 10^4^ CFU/mL. This resulted in the transfer of ~3 x 10^5^ or ~3 x 10^2^ CFU/mL to the hub and ~3 x 10^5^ or ~3 x 10^2^ CFU/mL to the peripheral leaves, for the high and low migration rates, respectively.

##### IN>OUT regime

60 μL of subpopulation 1, 3, and 4 (MIX) were mixed in 20 μL of fresh culture media with 20 ng/mL Ciprofloxacin and serially diluted to reach ~9 x 10^7^ or ~9 x 10^4^ CFU/mL. Also, 20μL of subpopulation 2 (HUB) was mixed in 180 μL of fresh culture media with 20 ng/mL Ciprofloxacin and serially diluted to reach ~10^7^ or ~10^4^ CFU/mL. This resulted in the transfer of ~9 x 10^5^ or ~9 x 10^2^ CFU/mL to the hub and ~10^5^ or ~10^2^ CFU/mL to the peripheral leaves, for the high and low migration rates, respectively.

#### Low population size

##### Well-mixed

20 μL of subpopulations 1, 2, 3, and 4 (MIX) were mixed in 120 μL of fresh culture media with 20 ng/mL Ciprofloxacin and serially diluted to reach ~4 x 10^5^ CFU/mL. Then, 15 μL of the diluted MIX was added to 1.5 mL fresh media along with 15 μL of 1:100 diluted the previous day’s culture. This resulted in the transfer of ~4 x 10^3^ CFU/mL for the migrant and ~10^5^ CFU/mL residents (“self-inoculation”).

##### IN>OUT Star

60 μL of subpopulations 1, 3, and 4 (MIX) were mixed in 20 μL of fresh culture media with 20 ng/mL Ciprofloxacin and serially diluted to reach ~9 x 10^5^ CFU/mL. Also, 20 μL of subpopulation 2 (HUB) was mixed in 180 μL of fresh culture media with 20 ng/mL Ciprofloxacin and serially diluted to reach ~10^5^ CFU/mL. This resulted in the transfer of ~9 x 10^3^ CFU/mL to the hub and ~10^3^ CFU/mL to the peripheral leaves, respectively. Every subpopulation also received 15 μL of 1:100 diluted the previous day’s culture from itself which resulted in the transfer of ~10^5^ CFU/mL residents (“self-inoculation”).

**Supplementary Figure 5:**
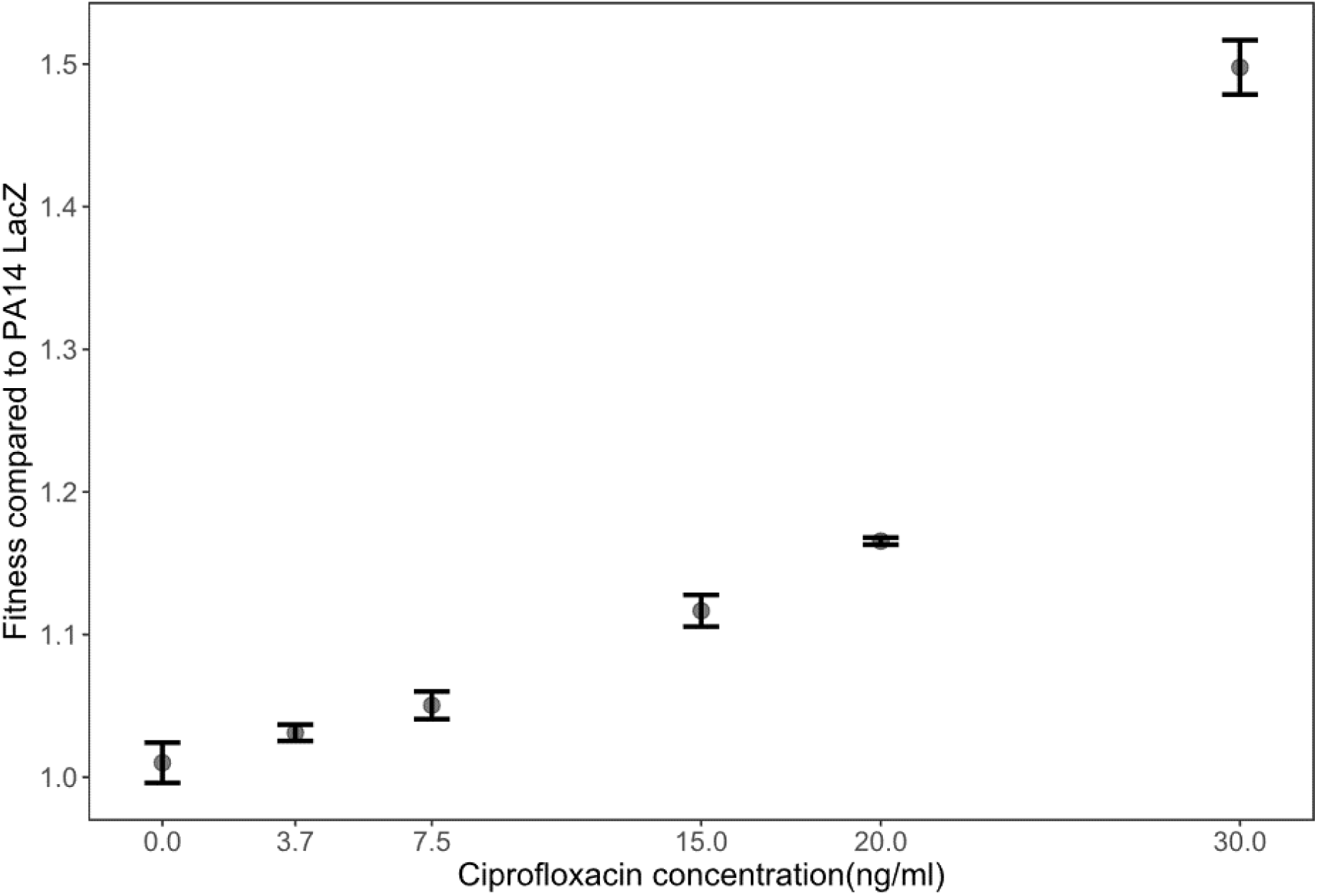
Competitive advantage of *PA14-gyrA* in head-to-head competitions (~ 50:50) with *PA14-LacZ* at different sub-inhibitory Ciprofloxacin concentrations. Relative fitness of three biological replicate competitions are shown.

**Supplementary Figure 6:**
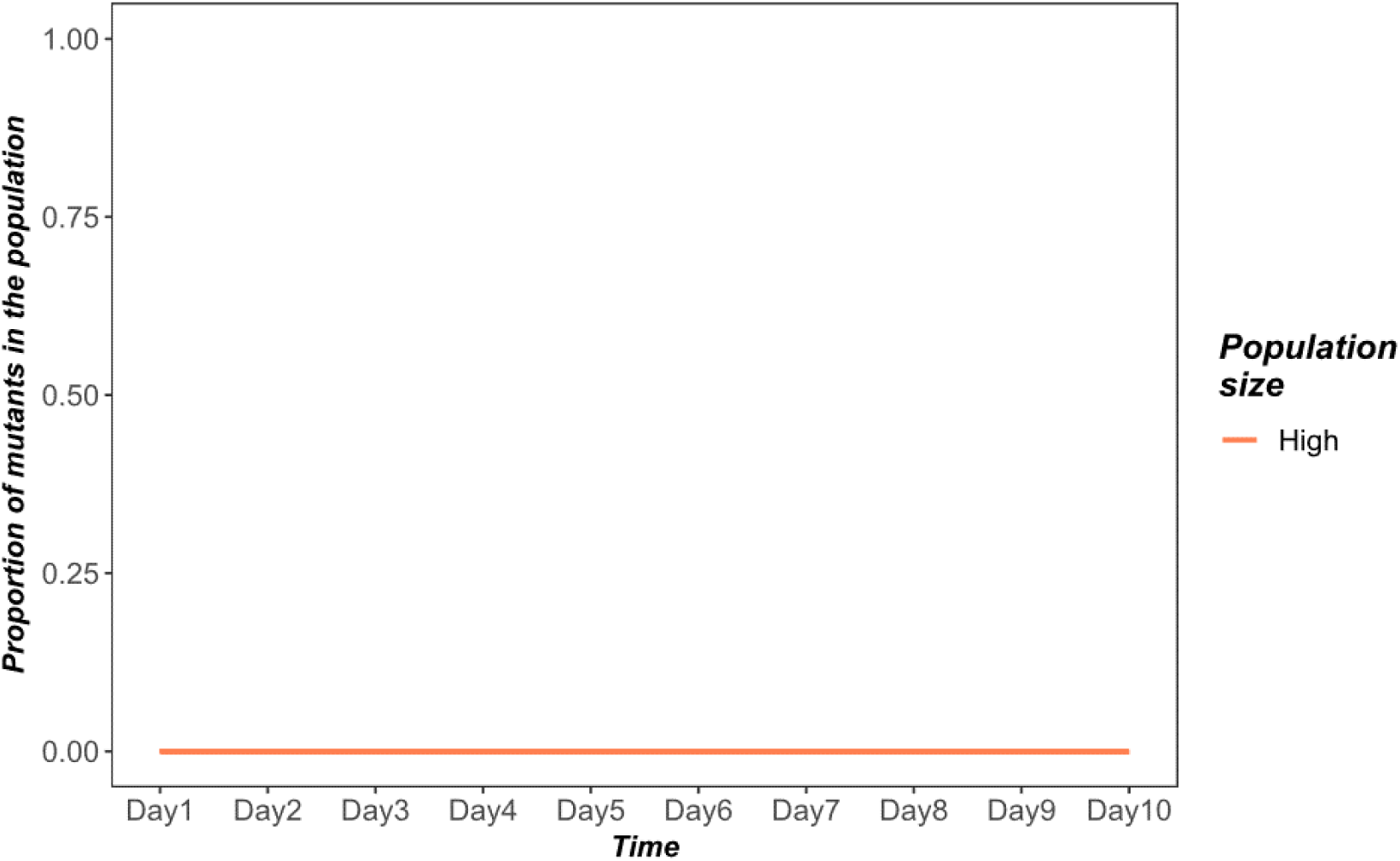
Frequency of spontaneous ciprofloxacin resistant (≥ 1μg/ml) mutants derived from the sensitive wild type strain, PA14-*LacZ*, over 10 days of experimental evolution in 20ng/ml Ciprofloxacin. A proportion of zero means there were no detectable colonies capable of growing above the MIC of the cip^R^ strain *PA14-gyrA* (T83I) used in our experiments.

**Supplementary figure 7:**
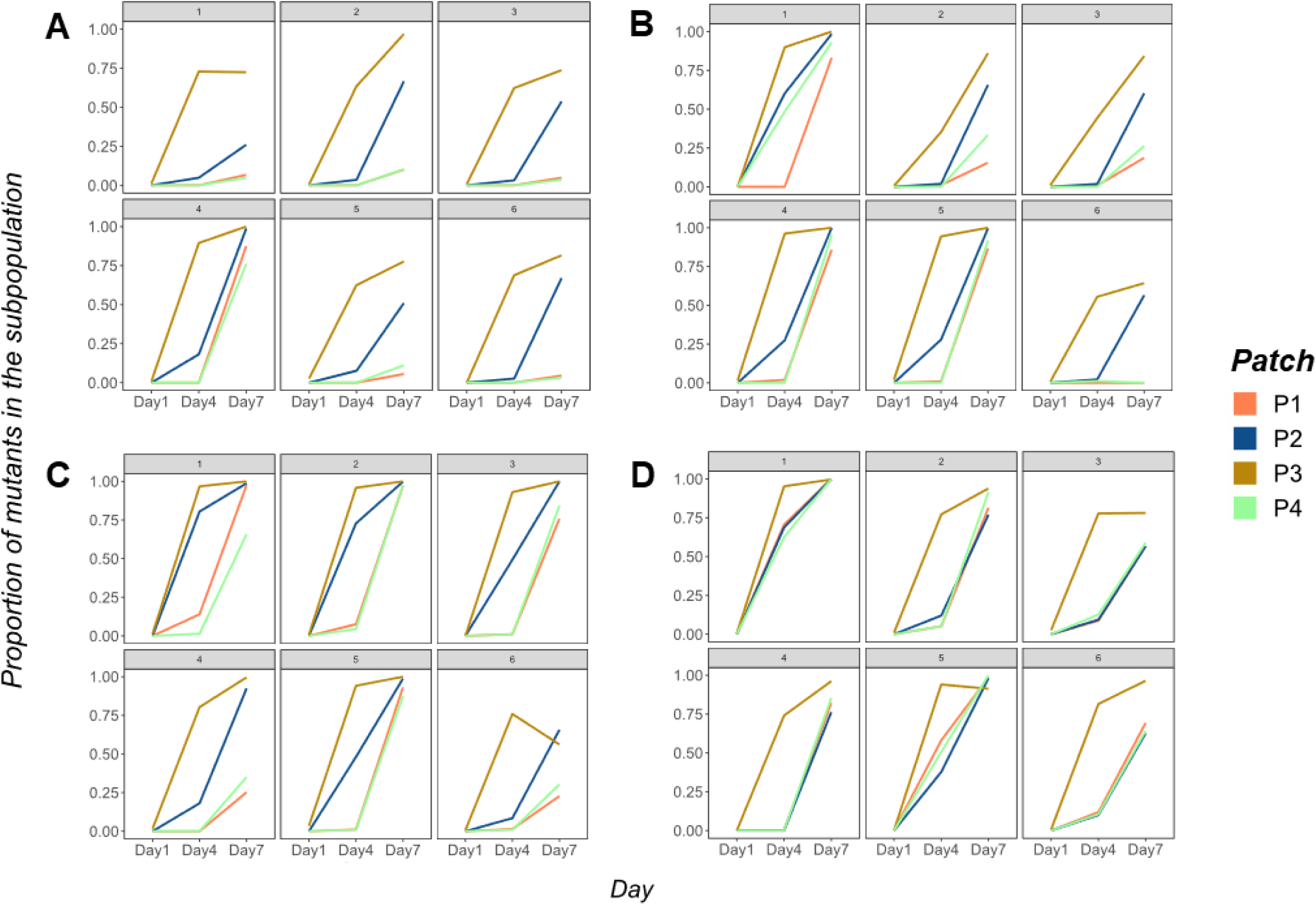
The proportion of the cip^R^ mutants in the constituent subpopulations of each metapopulation propagated by (A) IN=OUT, (B) OUT>IN, (C) IN>OUT and (D) well-mixed networks under high (10^5^ individuals) migration. Fixation dynamics for each replicate metapopulation is shown as panels. Subpopulation nomenclature: P3 = node of introduction of the cipR mutant, P2 = hub and P1 and P4 = rest of the peripheral leaves.

## Acknowledgement

We thank Caroline de Ligneris for her help around the lab. This work was supported by a Natural Sciences and Engineering Research Council (NSERC) Discovery Grant to RK.

## Author Contribution

R.K. conceptualized the project and provided guidance on lab techniques and analyses. P.P.C. performed the experiments and analyzed the data. L.R.N. performed the simulations and provided conceptual inputs. P.P.C, L.R.N. and R.K. wrote the manuscript.

## Materials & correspondence

All correspondence and requests for materials should be directed to Rees Kassen ‥

